# RodA Promotes Intestinal Colonization by Group B *Streptococcus*

**DOI:** 10.1101/2025.10.27.684869

**Authors:** Michelle J. Vaz, Sanjana Sankaran, Molly E. Sharp, Emily Dembinski, Adam J. Ratner

## Abstract

**Background:** Group B *Streptococcus* (GBS) intestinal colonization is critical for the pathogenesis of late-onset (LO) disease in infants. Using a murine model, we explore the role of *rodA*, which encodes a peptidoglycan polymerase RodA, belonging to the Shape, Elongation, Division, and Sporulation (SEDS) family that participates in peptidoglycan synthesis and maintenance of cell wall integrity.

**Methods:** We investigated the contribution of *rodA* to GBS gastrointestinal (GI) colonization using a wild-type strain (A909 WT) and an isogenic in-frame deletion mutant of *rodA* (A909Δ*rodA*). Morphological differences between the two strains were examined by transmission electron microscopy (TEM), and the contribution of *rodA* to GI colonization was assessed in a murine model through monocolonization and cocolonization experiments. We evaluated the growth of the mutant strain under intestinal physiological stress conditions and characterized its interactions with host epithelial cells in vitro.

**Results:** A909Δ*rodA* showed a unique chaining/aggregation phenotype compared to the A909 WT strain, with the presence of capsule confirmed via TEM and immunoblotting. In murine cocolonization experiments, A909 WT outcompeted A909Δ*rodA*; however, monocolonization experiments exhibited comparable colonization and bacterial burden across the GI tract. In vitro experiments revealed impaired growth in bile and an increase in adhesion to intestinal epithelial cells by the Δ*rodA* mutant.

**Conclusion(s):** *rodA* plays a role in GBS intestinal colonization. Deletion of *rodA* increases sensitivity to gastrointestinal stressors in vitro and causes a pronounced defect in competition in vivo, suggesting that the presence of *rodA* increases the fitness of GBS in the gut.

## Introduction

*Streptococcus agalactiae* (Group B *Streptococcus* [GBS]) is a leading etiology of neonatal infection (1). Implementation of intrapartum antibiotic prophylaxis has markedly reduced the incidence of early onset (EO) infection (occurring ≤ 7 days of life) (2, 3). However, rates of maternal rectovaginal colonization persist, and the incidence of late onset (LO) infection (occurring ≥7 days of life) remains unchanged, making LO disease the predominant form of invasive GBS infection in infants (4, 5). Recent work has shown the incidence of LO disease amongst a cohort of extremely preterm infants (< 28 weeks) as 8.47/1000 infants, nearly 30 times the national incidence of all neonates (6). LO disease has an overall case fatality rate of 4-12%, with the rates in preterm infants being twice that of term infants (3, 7). The sequelae of LO disease include significant neurodevelopmental impairment in survivors, adding to the morbidity burden (2, 8–11).

Prior studies have demonstrated that gastrointestinal (GI) colonization by GBS in infants is a fundamental precursor for LO infection (12, 13). Understanding how GBS establishes itself in the bacterial community of the intestinal tract can uncover mechanisms that contribute to the pathogenesis of LO disease. Our group recently showed the contribution of capsular polysaccharide, an important GBS virulence factor, to GI fitness and colonization in a murine model of GBS intestinal colonization (14). Hence, robust animal models facilitate examination of bacterial factors that augment or inhibit GI colonization.

In this report, we explore the role of GBS *rodA*, a gene that encodes the peptidoglycan polymerase RodA, which belongs to the SEDS family of proteins. RodA, through its role as a glycosyltransferase and in conjunction with penicillin-binding proteins (PBPs), is responsible for peptidoglycan synthesis, which is vital for cell wall integrity and responsible for bacterial survival under environmental stress (15, 16). It is known to contribute to bile salt resistance by GBS (17). In this report, we generated an in-frame deletion of *rodA* in the GBS A909 background using CRISPR-Cas12a mutagenesis and studied its impact on the cell wall and capsule. We analyzed its response to intestinal physiological stressors and determined its role in a postnatal murine model of GI colonization.

## Methods

### Bacterial Strains and Growth Conditions

GBS strain A909 (serotype Ia, ST7) and its derivatives were grown to stationary phase at 37°C in tryptic soy (TS) broth supplemented with 5 μg/ml erythromycin as needed for selection. *Escherichia coli* strain DH5α was grown at 37°C, shaking in Luria-Bertani (LB) medium supplemented with 300 μg/ml erythromycin when needed for selection.

### Construction of the GBS *rodA* deletion mutant and its complemented control strain

The *rodA* deletion mutant was created via a Cas12a-based platform as previously described (18). Briefly, a Cas12a-compatible genomic target site in the *rodA* gene was identified using the CRISPick server hosted by the Broad Institute (19). After selecting a target sequence, we designed ssDNA oligonucleotides and annealed them to make a dsDNA protospacer. The protospacer was then cloned into pGBSedit that had been linearized with Esp3I. The pGBSedit plasmid encodes *cas12a*, regulated by a P*_xyl/tet_* promoter, that responds to the presence of anhydrotetracycline (aTC). Primer pairs were used to amplify upstream and downstream homology arms flanking *rodA*. These fragments were cloned into pGBSedit linearized with XhoI using Gibson assembly. Following transformation into *E. coli*, plasmids were verified via PCR and whole-plasmid sequencing. Sequenced plasmids were transformed into electrocompetent GBS and successful transformants were confirmed by colony PCR (cPCR). Transformants were then subcultured in 10 mL TS + erythromycin (5 μg/mL) and allowed to grow at 37 °C for 6–8 hours. Subsequently, the cultures were subjected to aTC induction (500 ng/ml), resulting in strong selection against WT strains and for homology-driven recombination mutants (20). cPCR was used once again to confirm deletion mutants. Once confirmed, colonies were inoculated into 40 mL TS broth and grown overnight at 37°C. After a second passage of overnight growth in 40 mL of TS broth, 1:10,000 and 1:100,000 dilutions were plated on TS agar plates. The next day colonies were dual patched onto TS and TS + erythromycin (5 μg/mL) agar plates to detect cured clones. All constructs were confirmed by PCR and sequencing. We complemented the *rodA* gene deletion by using the pGBScomp High plasmid (A909Δ*rodA+*pGBScomp High::*rodA)* (18). Briefly, pGBScomp High plasmid was linearized via SalI restriction digest, and the *rodA* gene was PCR amplified. The two products were Gibson assembled and transformed into *E. coli* followed by confirmation by sequencing. Electrocompetent GBS was transformed with the plasmid and confirmed using cPCR. Primers used in the construction of the mutant and complemented strain are listed in **(Supplemental Table 1).** The transposon *rodA* mutant (A909 *rodA::Himar1*) was used from a large-scale GBS indexed library of *Himar1* mini-transposon mutant strains, containing interruptions of 878 genes and 253 intergenic regions, each with an erythromycin resistance cassette (21).

### Microscopy and Staining

#### Gram Stain

WT A909 and A909Δ*rodA* strains were grown overnight in TS broth at 37 °C. Cells were harvested by centrifugation (4,000 rpm, 10 min), washed once with Dulbecco’s phosphate-buffered saline (DPBS), and resuspended in DPBS. Smears were prepared on clean glass slides, heat-fixed, and subjected to Gram staining using the standard four-step protocol: crystal violet (1 min), Gram’s iodine (1 min), decolorization with 95% ethanol (10–15 s), and counterstaining with safranin (30 s), as previously described (22). Slides were rinsed gently with distilled water between each step and air-dried. Stained preparations were examined by light microscopy (1000×, oil immersion). Brightfield images were acquired using a Zeiss Axio Observer microscope equipped with a 63×/1.4 oil immersion objective.

#### Colony immunoblotting

Overnight cultures of WT A909, A909Δ*rodA* and A909Δ*cpsE* (acapsular control) were spread on CHROMagar plates. Colonies were counted to identify plates with 20–200 colonies for subsequent immunoblotting. Plates were briefly overlaid with nitrocellulose membranes (Amersham) to allow adherence of GBS material to the membrane. Membranes were blocked in 3% bovine serum albumin (BSA) in DPBS (blocking solution) for 1 hour. Blots were then transferred to type Ia *Streptococcus* group B type antisera (Statens Serum Institut) and incubated for 30 minutes with gentle shaking. Type Ia antiserum was diluted 1:2,000 in the blocking solution. Blots were then washed three times in DPBS. Blots were incubated in horseradish peroxidase (HRP)-conjugated goat anti-rabbit secondary antibody (Pierce) diluted 1:1,000 in blocking solution for 2 hours. Blots were washed three times in DPBS and stained using a 3,3′-diaminobenzidine tetrahydrochloride substrate kit (Abcam).

#### Electron microscopy

Overnight cultures were pelleted for 5 min at 4,000 rpm. Samples were processed and imaged by the NYU Microscopy Laboratory as previously described with modifications (23). In brief, bacteria were resuspended in 1 mL LRR (Lysine-Ruthenium Red-Osmium) fixative solution (1 mL fixation solution per sample: 0.5 mL 0.15% ruthenium red, 125 μL 16% formaldehyde, 100 μL 25% glutaraldehyde solution, 0.0155 g lysine acetate; fill with distilled water to 1 mL). The cells were fixed on ice for 20 min, washed two times with 0.1M sodium cacodylate buffer (0.1M cacodylate, 0.01M CaCl_2_, 0.01M MgCl_2_, 0.09M sucrose, pH 6.9) containing 0.075% ruthenium red and fixed in the fixation solution without lysine acetate for 2 hrs on ice. After washing three times with 0.1M sodium cacodylate buffer containing 0.075% ruthenium red for 5 min each, the cells were postfixed in 1% osmium tetroxide (OsO_4_) with 0.075% ruthenium red in 0.1M sodium cacodylate buffer for 1h on ice, washed three times with 0.1M sodium cacodylate buffer with 0.075% ruthenium red for 5 min each, then embedded with 2% agar and *en block* stained with 0.5% uranyl acetate aqueous solution overnight at 4°C. The samples were washed with distilled water, dehydrated in a graded ethanol series (30%: 50%: 75%, 85%: 90%:100%:100%) on ice, infiltrated and embedded with LR White resin (Electron Microscopy Sciences, PA) at 55°C. Thin sections were cut using Leica UC6 ultramicrotome, collected on 200 mesh copper grids (Ted, Pella Inc. CA), and stained with 4% aqueous uranyl acetate for 5 minutes. Stained grids were examined under JEOL1400 Flash transmission electron microscope (Japan) and photographed with a Gatan Rio 16 camera (Gatan Inc. Pleasanton, CA). **GI colonization model.** All experiments were performed in accordance with the NYU Grossman School of Medicine’s Institutional Animal Care and Use Committee (IACUC). These experiments were conducted as previously described (24). Adult male and female C57BL/6J mice (8–12 weeks) were purchased from Jackson Laboratories (Bar Harbor, Maine), given at least 3 days to acclimate to the local facility, and mated as pairs and trios. Dams were monitored for the birth of litters, and animals aged 12–14 days (preweaning) were used for colonization. Two cohorts were used for cocolonization: one for cocolonization with A909 WT and A909Δ*rodA* strains and the other using A909 WT and A909 *rodA::Himar1*. Bacterial cultures were grown, centrifuged and resuspended in sterile DPBS, and a 1:1 mixture was made of the resuspended strains to be competed. Animals were orally fed using a sterile feeding tube with 10^8^ colony forming units (CFUs) of GBS resuspended in 50 µL of DPBS. Animals remained housed with their biological dams and were monitored daily for signs of illness and mortality. At predetermined time points (7- and 14-days post-infection), animals were euthanized, and the GI tract was harvested. Each portion (small intestine, cecum, and colon) was homogenized. Serial dilutions were plated on CHROMagar StrepB for the enumeration of GBS CFUs. In addition, dilutions were spread on CHROMagar plates, with and without antibiotic selection (erythromycin), for differentiation between A909 WT and A909 *rodA::Himar1*. A909 WT and A909Δ*rodA* were differentiated by picking individual colonies for screening using cPCR. For the monocolonization model, bacterial cultures were grown overnight to stationary phase and the optical densities of the two strains were normalized. Cultures were then centrifuged and resuspended in sterile DPBS to a final concentration of 10^9^ CFU/mL. Animals were orally fed using a sterile feeding tube with 10^8^ CFU of GBS resuspended in 50 µL of DPBS and remained with their biologic, non-colonized dam. The animals were monitored daily for signs of illness and mortality. At predetermined time points (3-, 7-, and 14-days post-infection), animals were euthanized, and the GI tract was harvested. Each portion (small intestine, cecum, and colon) was homogenized, and serial dilutions were plated for enumeration of GBS CFUs on CHROMagar.

#### Colony differentiation using cPCR

Bacterial colonies recovered from the cocolonization experiment were screened for the presence or absence of *rodA* using cPCR. We selected ∼20 colonies per site per mouse (∼100 colonies per site), from organ dilutions that had been spread on CHROMagar plates. Selected colonies were patched on TS plates overnight, then resuspended in 20 µL of PBS, of which 2 µL was used as a template in a 25 µL PCR reaction containing OneTaq 2xMM, forward and reverse check primers **(Supplementary Table 1)** specific for the flanking region of the *rodA* gene and RNAse-free water. PCR was conducted using an annealing temperature of 54°C and 2-minute extension. Products were run on 1% agarose gel. Amplicon length was compared to A909 WT and A909Δ*rodA* controls to confirm genotype. Competitive index for cocolonization experiments was calculated as: (CFU strain 1 recovered/CFU strain 1 inoculated)/(CFU strain 2 recovered/CFU strain 2 inoculated) and log-transformed for calculation of geometric mean and 95% confidence interval (CI) for each GI site.

#### Growth rate analysis under physiological stressors

To investigate bacterial growth rate under acidic pH, bile, and lysozyme conditions, overnight cultures were assayed as described previously (17, 25, 26). TS broth was adjusted to pH 6.0 and 5.0 using sterile 1 N HCl. TS broth was prepared and autoclaved, cooled to room temperature (or 37 °C for temperature-matched readings), and stirred aseptically while titrating with sterile 1 N HCl. pH was monitored with a calibrated pH meter until 6.00 ± 0.05 and 5.00 ± 0.05 was reached. Media were re-checked after 10 min equilibration, dispensed, and used the same day. Control medium (pH 7.0) was handled identically without acid addition. Bile concentrations (5 mg/ml and 10 mg/ml) from a stock concentration of 200 mg/ml and lysozyme concentration of 14 mg/ml from a stock concentration of 100 mg/ml (made in DPBS) were prepared in TSB and sterilized prior use. Lysozyme concentration was determined through preliminary testing of multiple concentrations, with the final concentration selected to yield ∼50% killing in the WT strain (LD_50_) (**Supplemental Figure S1)**. Overnight cultures were diluted to an OD₆₀₀ of 0.05 in TS broth, 150 µL of each culture was added to a 96-well plate in three technical replicates. Media-only wells served as negative controls, and all readings were normalized to the negative control. Growth curve assays were performed using the BioTek LogPhase 600 Microbiology Reader for 24 hours at 37 °C, with readings taken every 20 minutes.

#### Biofilm Assay

Biofilm assays were performed as previously described (27). Briefly, overnight bacterial cultures grown in TSB were diluted to an OD₆₀₀ of 0.05 in Todd Hewitt Broth (THB) with 1% glucose, incubated for 24 hours at 37°C with 5% CO_2_ in 12 wells, flat bottom tissue culture plates. Staining was performed the next day using 0.5% crystal violet, solubilized with 30% glacial acetic acid and 100uL of samples were transferred to 96 well cell culture plates for readings. Spectra Max M3 was used to obtain readings at OD_540_. Biofilms under stress conditions were tested by diluting bacteria in THB 1% glucose with the relevant stress media as described above.

#### Tissue culture assays

The human intestinal Caco-2 cell line (ATCC HTB-37; colorectal adenocarcinoma) was seeded in 24-well plates at a density of 1 x 10^4^ cells per well in Eagle’s Minimum Essential Medium (EMEM, ATCC 30-2003) supplemented with 20% (vol/vol) fetal bovine serum (FBS, HyClone Cytiva, Marlborough, MA) and 1% penicillin-streptomycin (Pen/Strep, 10,000 U/mL, Gibco, USA) and cultured at 37°C in a humidified 5% CO_2_ incubator (28). The T84 colonic epithelial cell line (ATCC CCL-248) were cultivated in Dulbecco’s Modified Eagle’s Medium and Ham F-12 medium (DMEM: F-12, ATCC 30-2006) supplemented with 10% FBS and 1% Pen/Strep (29). For adherence assays, bacterial strains were grown overnight in TS broth. On the day of the assay, bacterial strains were diluted to an OD₆₀₀ of 0.1 and grown until mid-log phase. Caco-2 and T84 cells were grown to confluence in 12-well and 24-well tissue culture-treated plates (Corning). One hour prior to the assay, cell culture media was replaced with antibiotic - and serum-free media. Cells were washed three times with DPBS prior to infection. Bacteria were washed three times with DPBS and then resuspended in DMEM: F-12 or EMEM, depending on the cell type and added to cells at a multiplicity of infection (MOI) of 10. Adhesion was assessed after 45 minutes. Cells were washed six times with DPBS, removed from plates with trypsin-EDTA, and lysed with 0.1% saponin (30). Lysates were serially diluted and plated on TS agar (TSA) plates to quantify the number of adherent CFU. Adherence was calculated as: (total CFU recovered/total CFU of original inoculum) × 100%. For invasion assays, bacterial strains were prepared in the same manner as for adherence. Following three washes, bacteria was added to each well, and plates were incubated for 150 minutes to allow invasion. Plates were then washed six times to remove non-adherent bacteria. Antibiotic-containing culture media was prepared by adding gentamicin and penicillin from prepared stocks of 50 mg/mL each with a final concentration of 100 µg/mL gentamicin and 5 µg/mL penicillin. Plates were incubated for an additional 2 hours with antibiotic media, after which invasion capacity was assessed using the same procedures described for adherence.

#### Statistical Analysis

Statistics were calculated using GraphPad Prism 10.5.0 software. Colonization rates between the A909 WT and A909Δ*rodA* strains were compared using Fisher’s exact test. Bacterial burden comparison between the A909 WT and A909Δ*rodA* strains at different GI tract sites were compared using two-way repeated measures ANOVA with Sidak’s multiple comparisons post-test. Biofilm formation under different stressors was evaluated using two-way ANOVA with the Tukey multiple comparisons test. Tissue culture adhesion and invasion assays were compared using Mann-Whitney U test.

## Results

### Structural characterization of GBS A909Δ*rodA*

To characterize the structural contribution of *rodA*, we generated an in-frame deletion mutant (A909Δ*rodA)* using the Cas12a mutagenesis system as previously described (18). Overnight growth showed pronounced sedimentation of the A909Δ*rodA* compared to the A909 WT strain and complemented A909Δ*rodA+*pGBScomp High::*rodA*, although no significant differences were noted in growth rate **(Supplemental Figure S2, Figure 4A)**. Gram staining revealed a typical streptococcal morphology, with chains of cocci in the WT strain whereas the A909Δ*rodA* strain displayed a marked aggregation phenotype, with cells forming dense clusters versus discrete chains **(Figure 1A).** Since GBS capsule is anchored to the peptidoglycan layer, we assessed capsule presence to determine if *rodA* played a role in its production (31). Colony immunoblotting using a capsule-specific antibody detected capsule on both the A909 WT and A909 Δ*rodA* strains in contrast to the A909Δ*cpsE,* which did produce capsule (**Figure 1B).** In addition, we confirmed the presence of capsule production in the A909Δ*rodA* strain by latex agglutination (**Figure 1C).** Transmission electron microscopy did not reveal obvious abnormalities in morphology or envelope appearance in the A909Δ*rodA* strain compared to the A909 WT strain **(Figure 1D)**. These findings suggest a phenotype with chaining and aggregation in the A909Δ*rodA* strain, without detectable changes in capsule expression or obvious structural defects.

**Figure 1.**
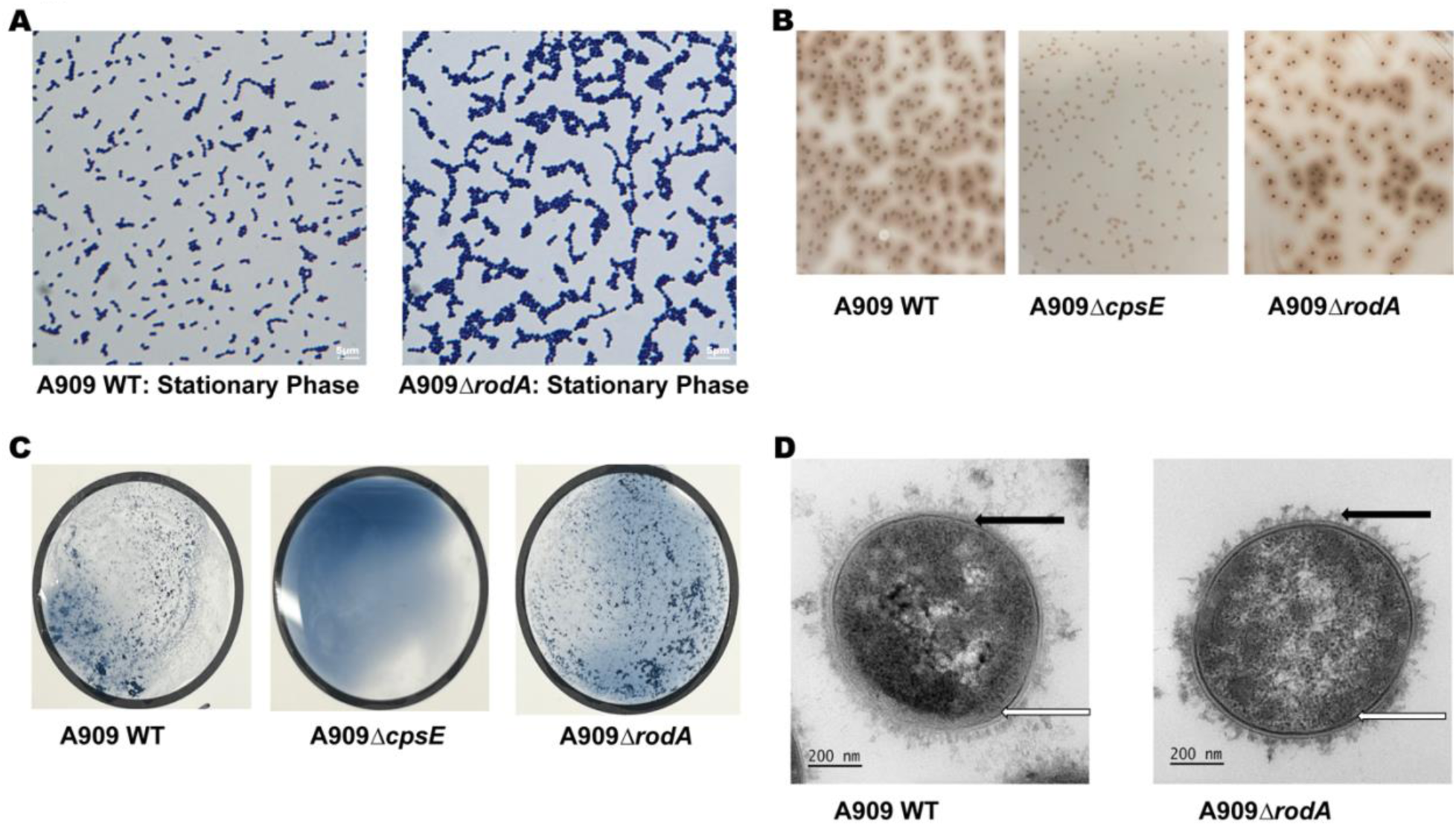
Structural characterization of WT and *rodA* deficient GBS. **A**. Gram-stained images of A909 WT and A909Δ*rodA* strains in stationary growth, taken at 63×original magnification. Scale bar = 5 µm. Experiments were performed twice independently, with representative images shown. **B.** Capsule production assessed by colony immunoblot using a serotype Ia-specific anti-capsular polysaccharide antibody in A909 WT, A909Δ*cpsE* (acapsular control) and A909Δ*rodA* strains. Positive: Halo around brown colonies; Negative: brown colonies only. Representative blot from two independent experiments is shown. **C.** Latex agglutination was used to confirm production of capsule. Type Ia antisera was used for A909 WT, A909 Δ*cpsE* (acapsular control) and A909Δ*rodA* strains. Positive: clumping noted; Negative: no clumping noted. Representative images from two independent experiments are shown. **D.** Transmission electron microscopy demonstrates capsule and cell wall in A909 WT and A909Δ*rodA* strains. Bacteria were chemically fixed by including lysine acetate during the glutaraldehyde and formaldehyde exposure. Samples were stained and postfixed using Ruthenium red and osmium and subsequently embedded in LR White. Black arrows mark the capsule, and white arrows mark the cell wall. Scale bar = 200 nm. Representative images from 10 fields per strains are shown.

### *rodA* provides a competitive advantage in GI colonization

To assess the contribution of *rodA* in GI colonization, we initially employed a *rodA* transposon mutant (A909 *rodA::Himar1*). In cocolonization experiments, we noted that A909 WT outcompeted A909 *rodA::Himar1,* with geometric mean competitive indices 6.62, 95% CI [0.85, 53.1] and 4.19, 95% CI [1.70, 10.35] in the small intestine; 7.67, 95% CI [1.69, 34.8] and 45.78, 95% CI [10.38, 201.7] in the cecum, and 7.5, 95% CI [1.68, 33.76] and 49.82, 95% CI [17.72, 140.1] in the colon, at 7 and 14 days respectively indicating a competitive advantage in vivo **(Supplemental Figure S3).** Based on these pilot findings, we used the in-frame deletion mutant (A909Δ*rodA*) to validate the in vivo phenotype. Preweaning mice (12–14 days) were orally infected in cocolonization experiments as described (24). Animals cocolonized with the A909 WT and A909Δ*rodA* strains were monitored for signs of illness and survival until 7 days post infection. There was 100% survival in this cohort at the experimental endpoint of 7 days, at which point animals were euthanized and the GI tract harvested **(Figure 2A)**. Colonies recovered from the small intestine, cecum, and colon were analyzed using cPCR to determine the ratio of A909 WT to A909Δ*rodA* **(Supplemental Figure 4)**. In the small intestine and cecum, the A909 WT strain outcompeted the A909Δ*rodA* strain with a geometric mean competitive index of 40, 95% CI [40, 40] at 7 days post-colonization, denoting no variation across replicates, as all tested colonies were A909 WT **(Figure 2B)**. In the colon, we calculated a geometric mean of 34.47, 95% CI [22.8, 52.11] **(Figure 2B)**. These findings suggest a sustained competitive advantage of the A909 WT strain over the A909Δ*rodA* strain throughout the GI tract.

**Figure 2.**
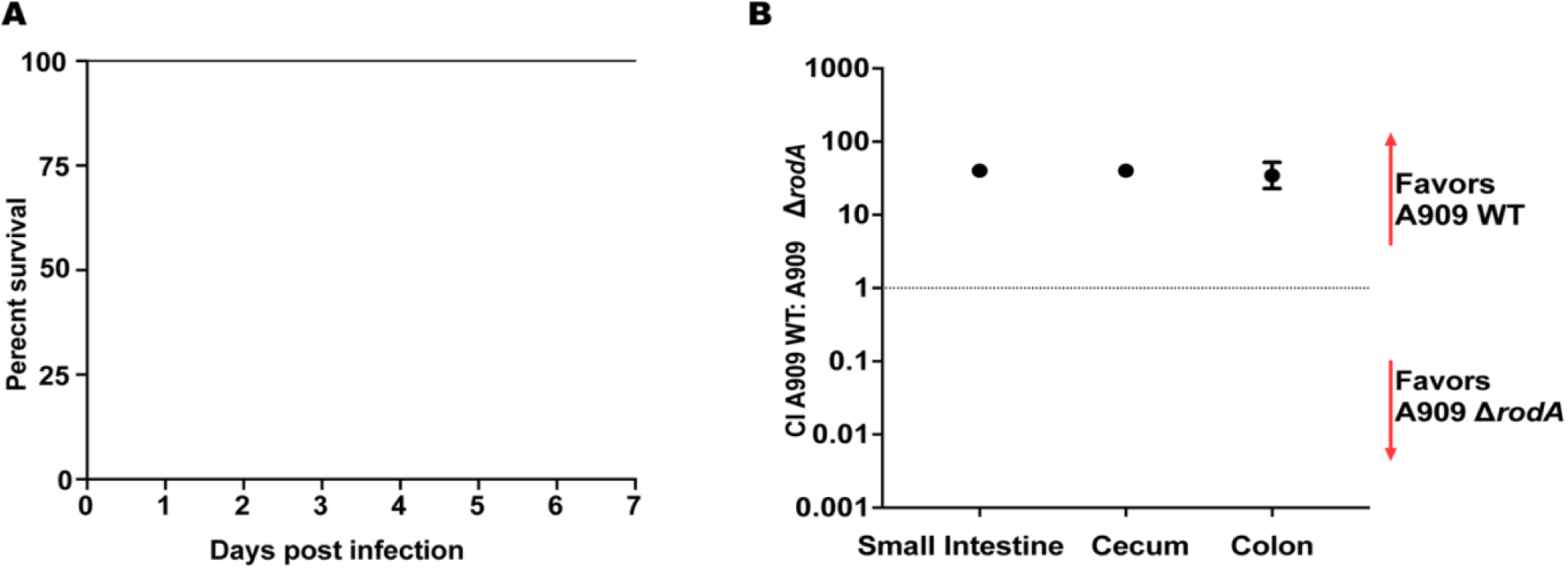
RodA confers a competitive advantage to GBS in an intestinal colonization competition model. Preweaning C57BL/6J mice were orally inoculated with a 1:1 mixture of A909 WT and A909Δ*rodA* in an experimental cohort (n = 5). **A.** Kaplan-Meier survival curve demonstrates 100% survival. **B.** The small intestine, cecum, and colon, were harvested at 7 days post inoculation. Approximately 100 colonies per GI site (∼20 colonies per mouse) were analyzed using cPCR to differentiate between A909 WT and A909Δ*rodA*. Data points represent geometric mean competition indices and error bars represent 95% confidence intervals.

### *rodA* is dispensable for intestinal colonization in the absence of competition

We next assessed the role of *rodA* in monocolonization. In longitudinal experiments across multiple cohorts, pre-weaning mice were orally infected with either the A909 WT strain or the A909Δ*rodA* strain. These animals were longitudinally monitored for signs or illness/mortality, with cohorts euthanized at days 3, 7 or 14. We noted equivalent bacterial colonization between the A909 WT and A909Δ*rodA* strains throughout the GI tract at all 3 time points **(Figure 3 A, C, E)**. On further assessment, we noted comparable bacterial burden throughout the GI tract, except in the colon at day 3 and day 7 post colonization, where the A909 WT strain had a higher median colonization density than the A909Δ*rodA* strain **(Figure 3 B, D, F)**. There was an overall decrease in bacterial burden at day 14 post infection, across strains **(Figure 3F)**. Taken together, we note that A909Δ*rodA* demonstrated adequate colonization efficacy in monocolonization experiments; however, it was outcompeted by the A909 WT strain in cocolonization. This suggests that *rodA* may not be strictly essential for establishment of intestinal colonization but may confer a selective advantage under competitive pressure.

**Figure 3.**
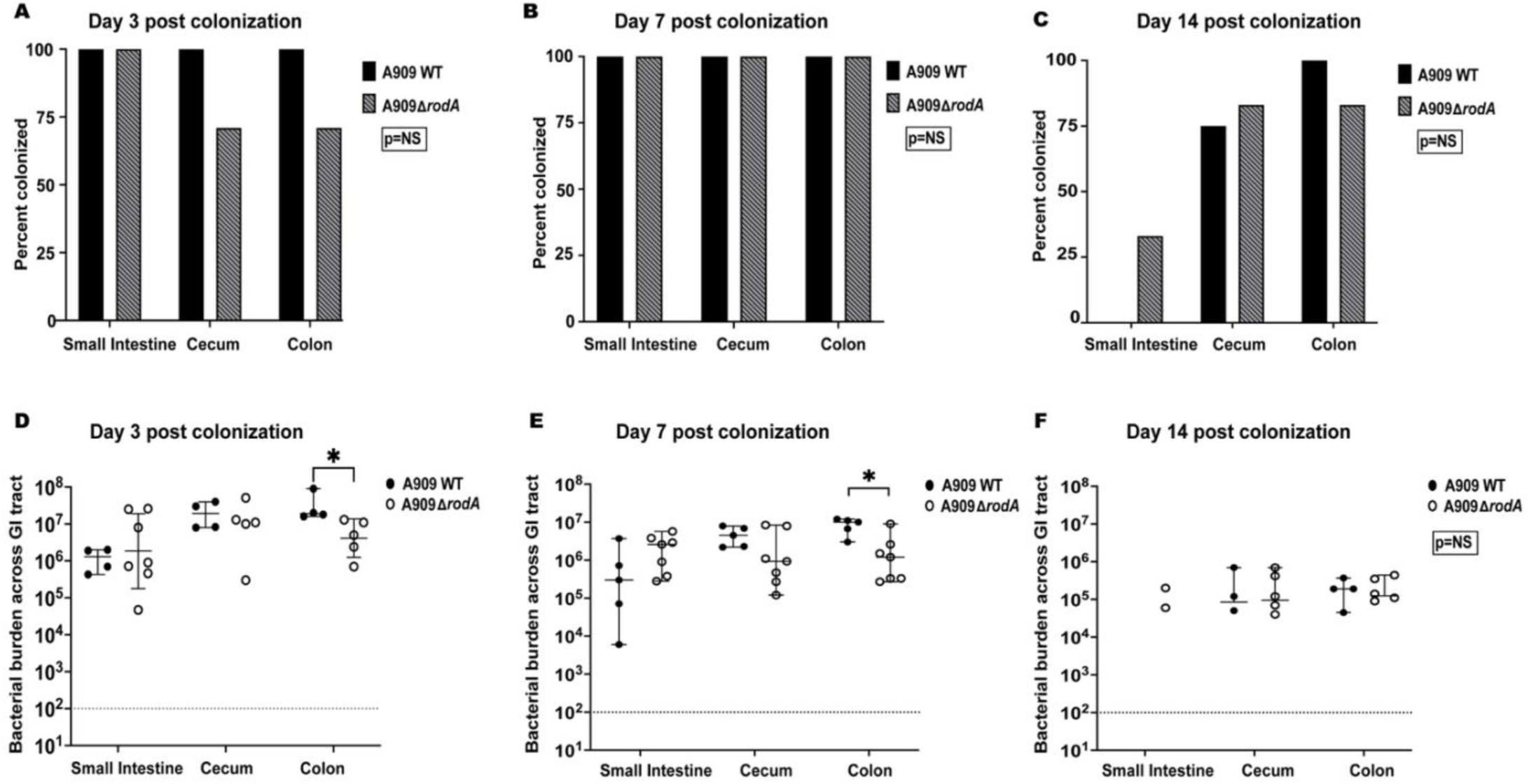
Comparable colonization by the A909Δ*rodA* mutant and A909 WT strains in monocolonization. Preweaning C57BL/6J mice were orally infected with either A909 WT or A909Δ*rodA* in six separate cohorts, and the small intestine, cecum, and colon, were harvested at predetermined time intervals. **A, B, C.** Percent colonization as determined by the presence of GBS on days 3, 7, and 14 respectively (P = not significant [NS]; two-sided Fisher’s exact test). **D, E, F.** Bacterial burden determined as GBS CFU/gram tissue on days 3, 7, and 14 respectively. Individual points represent biological replicates and lines showing median with interquartile range, (*, *p<0.01*; **, *p<0.001*; two-way repeated measures ANOVA with Sidak’s multiple comparisons post-test). Sample sizes: A909 WT=4, A909Δ*rodA*=7 **(Figure A, D)**, A909 WT = 5, A909Δ*rodA* = 7 **(Figure B, E)**, A909 WT=4, A909Δ*rodA* = 6 **(Figure C, F)**.

### Investigating the role of *rodA* in response to intestinal physiological stress

To further characterize the role of *rodA* in the GI tract, we explored bacterial growth in vitro in the presence of physiological gastrointestinal stress conditions. We monitored growth of A909 WT, A909Δ*rodA* and complemented strain (A909Δ*rodA*+pGBScomp High::*rodA*) in TSB with acidic pH in the range of 5.0-7.34, bile concentrations of 5 mg/mL and 10 mg/mL, and lysozyme (14 mg/ml). The growth of all three strains was unaffected by a pH of 6, however there was an overall decrease in growth rate and bacterial mass at a pH of 5 for all strains **(Figure 4B, C)**. The growth of A909Δ*rodA* was markedly inhibited in both bile salt concentrations, while the WT strain grew adequately, and the complemented strain exhibited partial restoration of growth at both bile concentrations **(Figure 4D, E)**. Short term lysozyme exposure resulted in ∼50% reduction in viability of A909 WT (**Supplemental Figure S1),** however we noted no significant difference in long term growth in both strains in the presence of lysozyme **(Figure 4F).** In all assays, the complemented strain showed a delay in entry into log phase, likely due to growth under antibiotic selection pressure. Together these findings suggest that *rodA* is not required for survival under acid stress and does not contribute to lysozyme resistance in long term assays, but it is required for optimal growth in the presence of bile salts.

**Figure 4.**
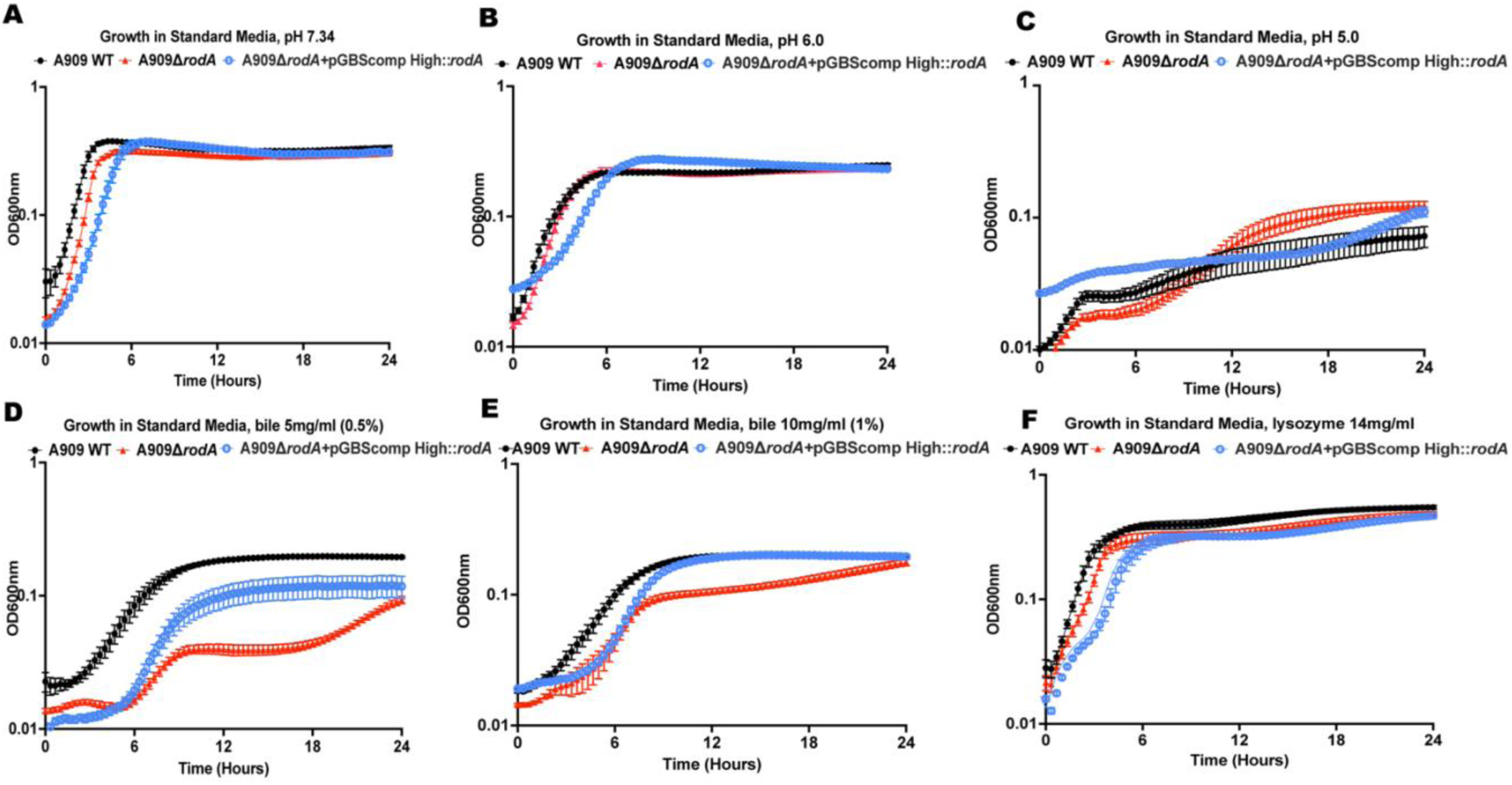
Exposure to intestinal physiological stressors reveals selective susceptibility of A909Δ*rodA*. Overnight cultures of A909 WT, A909Δ*rodA* and A909Δ*rodA+*pGBScomp High::*rodA* GBS strains were diluted to an OD value of 0.05 in Todd-Hewitt broth and incubated at 37°C. OD_600_ was measured at 20-minute intervals under different intestinal stressors. **A.** Standard media (pH 7.34), **B.** Standard media, pH 6.0, **C.** Standard media, pH 5.0, **D.** Standard media, bile 5 mg/ml (0.5%), **E**. Standard media, bile 10 mg/ml (1.0%), **F.** Standard media, lysozyme 14 mg/ml. Data from independent samples (n = 9) with three technical replicates shown, with lines indicating mean values ± standard errors of the mean (SEM).

### *rodA* influences bacterial surface contact to epithelial surfaces and modulates host interactions

As biofilm formation and adhesion to host epithelial surfaces are both important for gastrointestinal persistence, we next examined whether deletion of *rodA* altered these phenotypes. We assessed the formation of biofilm in A909 WT and A909Δ*rodA* strains after exposure to acidic media, lysozyme and bile. At pH 6.0, no difference was noted in biofilm production between the WT and mutant strains **(Figure 5A)**. At pH 5.0, the A909Δ*rodA* strain exhibited a non-significant decrease in biofilm production compared to its control (standard media, pH 7.34) **(Figure 5B)**. On exposure to lysozyme, we noted an increase in biofilm formation by the A909 WT strain that was not seen in the A909Δ*rodA* strain **(Figure 5C)**. The presence of bile salts decreased the formation of biofilm in both the WT and mutant strains when compared to their respective controls **(Figure 5D, E)**. To understand the surface associated behavior of *rodA* at epithelial barriers, we performed adherence and invasion assays using differentiated Caco-2 intestinal epithelial cells and T84 colonic epithelial cell lines, in an in vitro model. We evaluated if the adherence and/or invasion to epithelial cells by GBS depended on *rodA*. Adherence assays showed increased adherence of the A909Δ*rodA* strain in the Caco-2 and T84 cell lines compared to the A909 WT strain **(Figure 6A, B)**. In contrast, invasion assay results differed by cell type. In Caco-2 cell, there was no difference in invasion between the A909 WT and A909Δ*rodA* strains, whereas in T84 cells, there was an increase in invasion by the A909Δ*rodA* strain **(Figure 6C, D)**. These findings suggest that the loss of *rodA* alters biofilm responses to gastrointestinal stressors and increases epithelial adherence.

**Figure 5.**
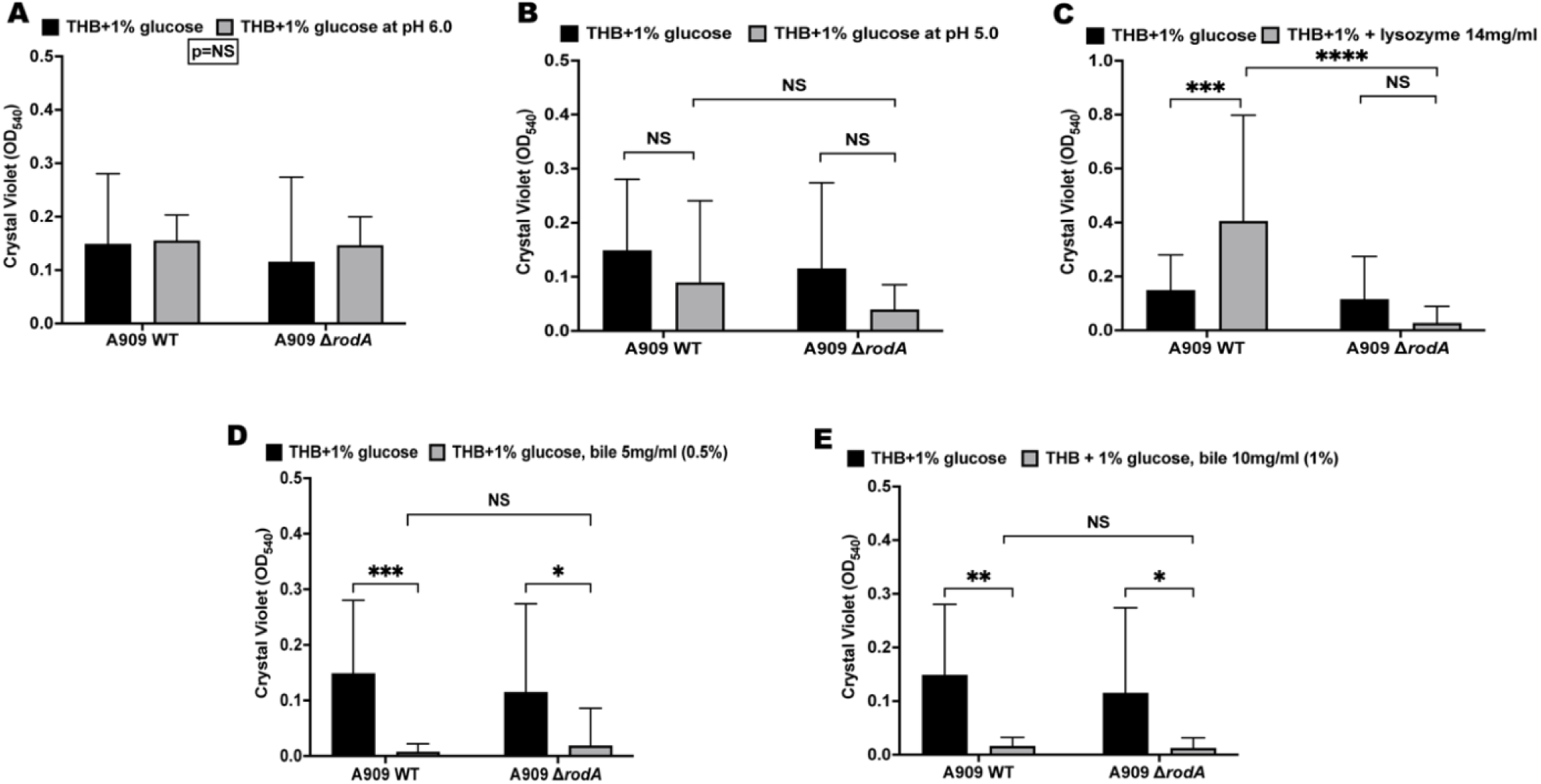
GBS *rodA* deficiency impacts biofilm formation under select stressor exposure. Biofilm formation of A909 WT and A909Δ*rodA* strains quantified by crystal violet uptake after exposure to **A.** THB, with 1% glucose at pH 6.0, **B.** THB, with 1% glucose at pH 5.0, **C.** THB, with 1% glucose, lysozyme 14 mg/ml, **D.** THB, with 1% glucose, bile 5 mg/ml, **E.** THB, with 1% glucose, bile 10 mg/ml. Data from independent samples (n = 9) with three technical replicates are shown, and lines represent mean values ± SDs. Data were analyzed using two-way ANOVA with the Tukey multiple comparisons test (*, *p<0.05*; **, *p<0.005*; \*\*\*\**p< 0.0001)*.

**Figure 6.**
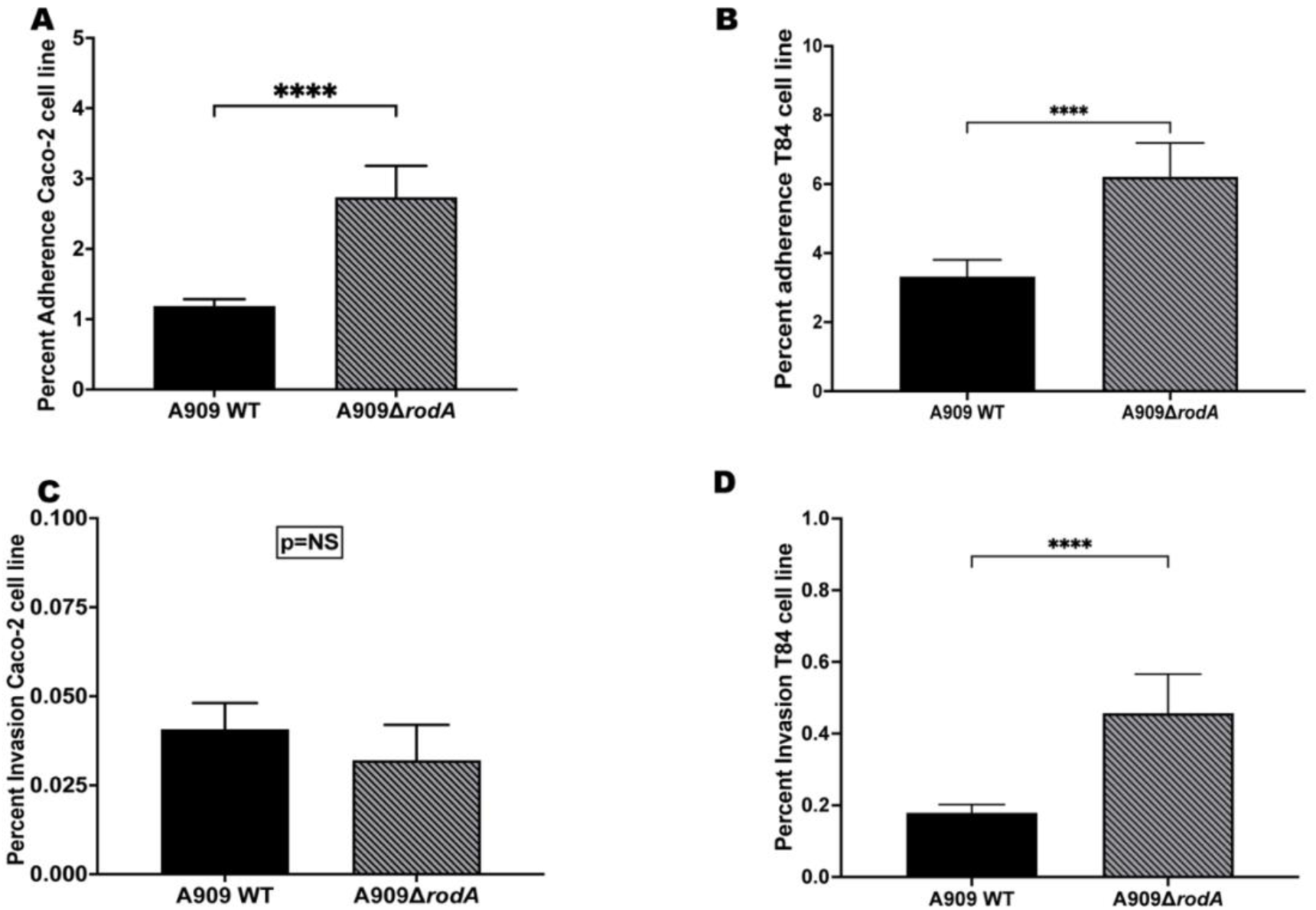
The absence of *rodA* impacts adherence and differentially affects invasion in intestinal epithelial cells. A, **C.** Adhesion assays were performed with Caco-2 and T84 intestinal cell lines respectively at an MOI of 10. **B, D.** Invasion assays were performed with Caco-2 and T84 intestinal cell lines respectively at an MOI 10. The percentage of adhered and invaded bacteria were calculated relative to the initial inoculum. All data are from independent experiments (n=9, each with 3 technical replicates). The error bars indicate the 95% confidence intervals of the means of the wells. Data were analyzed using the Mann-Whitney U test (****, *p < 0.0001)*.

## Discussion

The GBS cell wall is composed of a thick layer of peptidoglycan that anchors surface molecules and consists of an outermost boundary of capsular polysaccharide that serves as a major factor in colonization and pathogenesis (32–35). SEDS-family proteins, including RodA, are conserved transmembrane enzymes that are key meditators of peptidoglycan synthesis and cell wall integrity (36). RodA is a peptidoglycan polymerase that elongates glycan strands (16). It works with PBP’s, known to mediate peptide crosslinking, together playing a key morphogenetic role in bacterial cell wall synthesis (37, 38). Studies in several enteric pathogens illustrate that disruptions in glycan polymerization can have varying phenotypes depending on the organism and the environmental niche (37, 39–44). Laubacher et al. demonstrated that peptidoglycan stress activates Rcs envelope stress response in *E. coli*, leading to upregulation of capsule polysaccharide genes and contributing to increased intrinsic resistance in the presence of antibiotics (40). Prior reports have shown that the deletion of *rodA* leads to a loss of structure as seen in *Streptococcus pneumoniae*, *E. coli*, and *Bacillus subtilis* (39, 41, 44, 45). In contrast, our Δ*rodA* mutant showed no obvious changes to the bacterial shape by TEM and retained its capsular expression. This phenotypic difference from other bacteria may reflect species-specific differences in peptidoglycan synthesis or the relative contribution of RodA to overall cell wall integrity. In addition, as previously described in *Bacillus subtilis*, mechanisms such as alternative PG polymerases may compensate for the overall peptidoglycan integrity in GBS in the absence of RodA (37). Structural characterization also revealed a chaining/aggregating effect by A909Δ*rodA* on Gram staining. We propose that the aggregating and chaining phenomenon are due to subtle changes in peptidoglycan remodeling rather than a direct defect in division or capsule synthesis.

We noted increased adherence by the A909Δ*rodA* strain to both Caco-2 and T84 cell lines which may be driven by the pronounced chaining/aggregating phenotype. The difference in cell type specific responses to invasion, higher in the T84 than the Caco-2 cell line, may reflect specific characteristics of the host epithelial surface, including mucus production or barrier integrity. These results differ from *Salmonella*, where disruption of RodA-PBP complexes downregulates classical invasion factors such as flagella, which could imply that RodA modulates host interactions by physical rather than regulatory interactions (42). The biological significance of increased adherence with varied invasion in the mutant strain remain unclear and may reflect differences in morphological architecture of the epithelial cell lines, where Caco-2 cell line is known to show a distinct biochemical signature of mature small intestinal enterocytes whereas the T84 cell line retains its colonic differentiation (46).

In a model of GI colonization, we noted that the A909Δ*rodA* mutant displayed adequate colonization during monocolonization but was significantly outcompeted by the WT strain during cocolonization. We noted comparable bacterial burden across different sites of the GI tract, except the colon, in longitudinal experiments. This suggests that *rodA* is not essential for intestinal colonization but may confer an advantage under competitive pressure. This phenomenon of competitive interactions has been described in other enteric pathogens and is likely dependent on host, microbial, or nutritional pressures that become apparent during competition (47–49).

Exposure to intestinal physiological stressors like acid, bile and lysozyme showed that *rodA* is important for bile tolerance and biofilm adaptation to certain stressors. Jia et al. reported the conditional essentiality of the *rodA* gene to bile salt resistance by GBS (17). We hypothesize that the absence of *rodA* may impact the ability of the bacteria to polymerize glycan strands making the cell wall more permeable. This could explain the increase in bile susceptibility which disrupts the bacterial cell envelope by causing cell membrane leakage and denaturing proteins. Interestingly, the A909Δ*rodA* mutant showed greater susceptibility at 0.5% bile concentration compared to 1% bile concentration which was not seen in the A909 WT strain. This may indicate differential compensatory response of the A909Δ*rodA* mutant cell wall to bile and warrants further investigation. The reduced viability of GBS in short term assays but absence of long-term growth defect on lysozyme exposure, may reflect bacterial killing with subsequent recovery of surviving cells or obscuring of early CFU loss due to OD_600_ measurements (50). We also noted WT GBS increased biofilm formation in response to lysozyme, consistent with stress-induced biofilm formation described in other Gram-positive bacteria (51, 52). The absence of this response in the Δ*rodA* mutant suggests that RodA contributes to envelope remodeling under host-derived stress.

The context-dependent role of RodA uncovered in this report reflects its biological relevance under stress/competition as well as its conditional requirement for envelope integrity. RodA permits GBS to withstand physiological stressors, particularly bile, and to compete in the GI tract. It gives us a window into the dynamic microbial changes that occur in a natural setting during colonization persistence and/or clearance. This work highlights the importance of identifying other conditionally essential genes like *rodA* that confer a competitive advantage in the GI tract to inform strategies to disrupt GBS colonization.

## Author Notes

This work was presented in part at the Pediatric Academic Societies’ 2025 Annual Meeting.

