## Supplementary Figures 1-4 for "RodA Promotes Intestinal Colonization by Group B *Streptococcus*"

**Supplementary Figure 1.** Optimization of Lysozyme Concentration. Lysozyme concentrations were determined through preliminary testing of multiple concentrations (0 mg/ml, 12 mg/ml, 14 mg/ml and 20 mg/ml), with the final concentration of 14mg/mL selected for the main experiment based on optimal cell lysis activity of 50% at 1 hour.

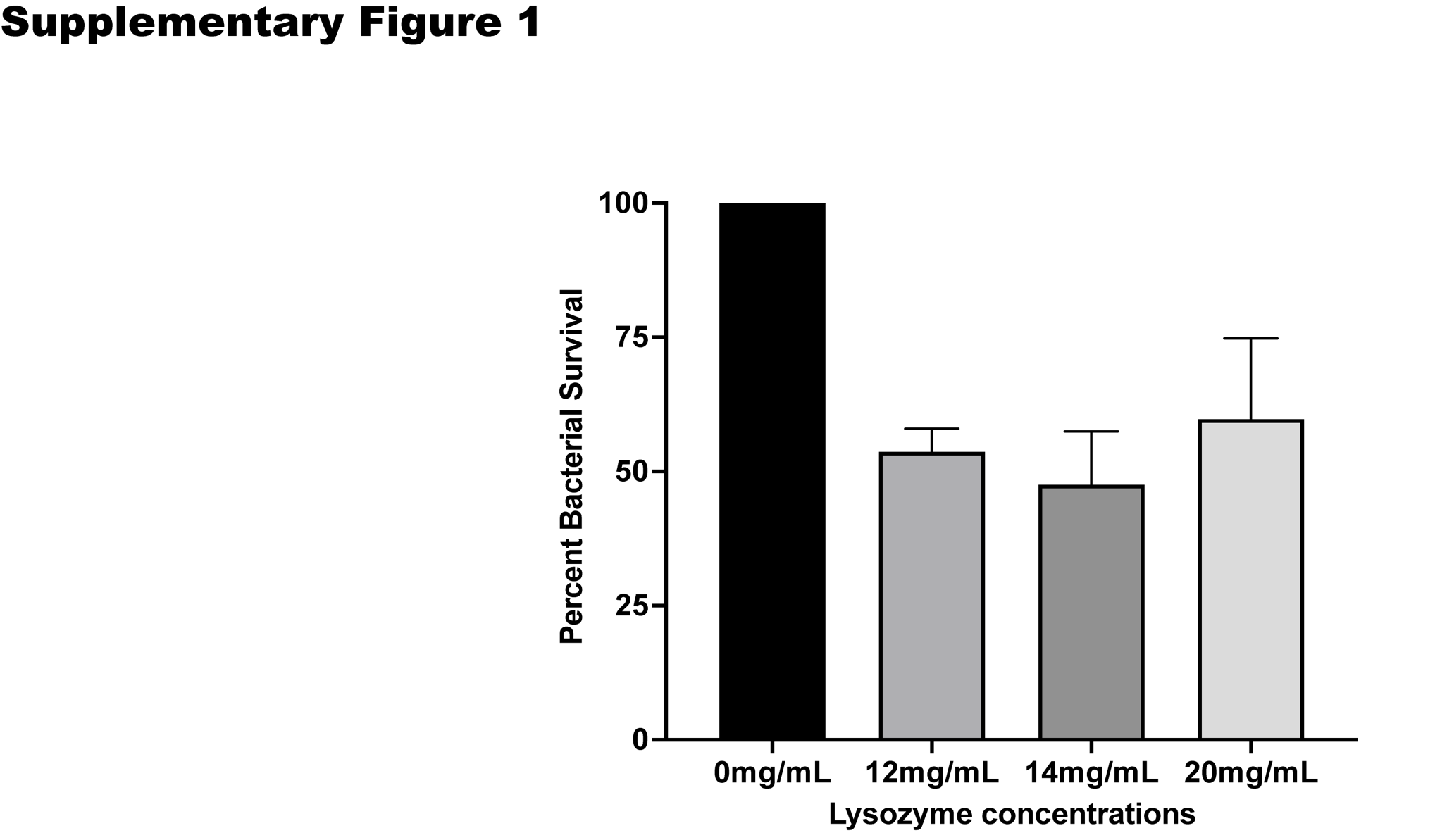

**Supplementary Figure 2.** Visualization of stationary-phase cultures grown for 18 hours at 37°C in Tryptic-Soy broth (TSB).

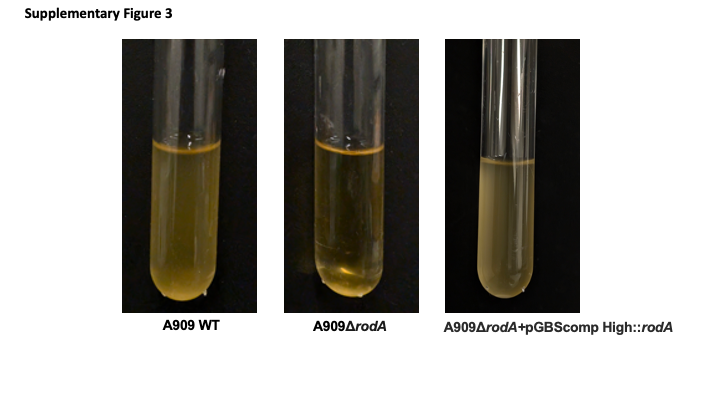

**Supplementary Figure 3.** Wild type (WT) A909 outcompetes *rodA* transposon insertion strain (A909 *rodA::Himar1*) in a cocolonization model. Preweaning C57BL/6J mice were orally inoculated with a 1:1 mixture of A909 WT and A909 *rodA::Himar1* in 2 separate experimental cohorts*.* The GI tract, small intestine, cecum, and colon were harvested at 7 days (n=8) and 14 days (n=6) post inoculation. Data points represent geometric mean competition indices and error bars represent 95% confidence intervals.

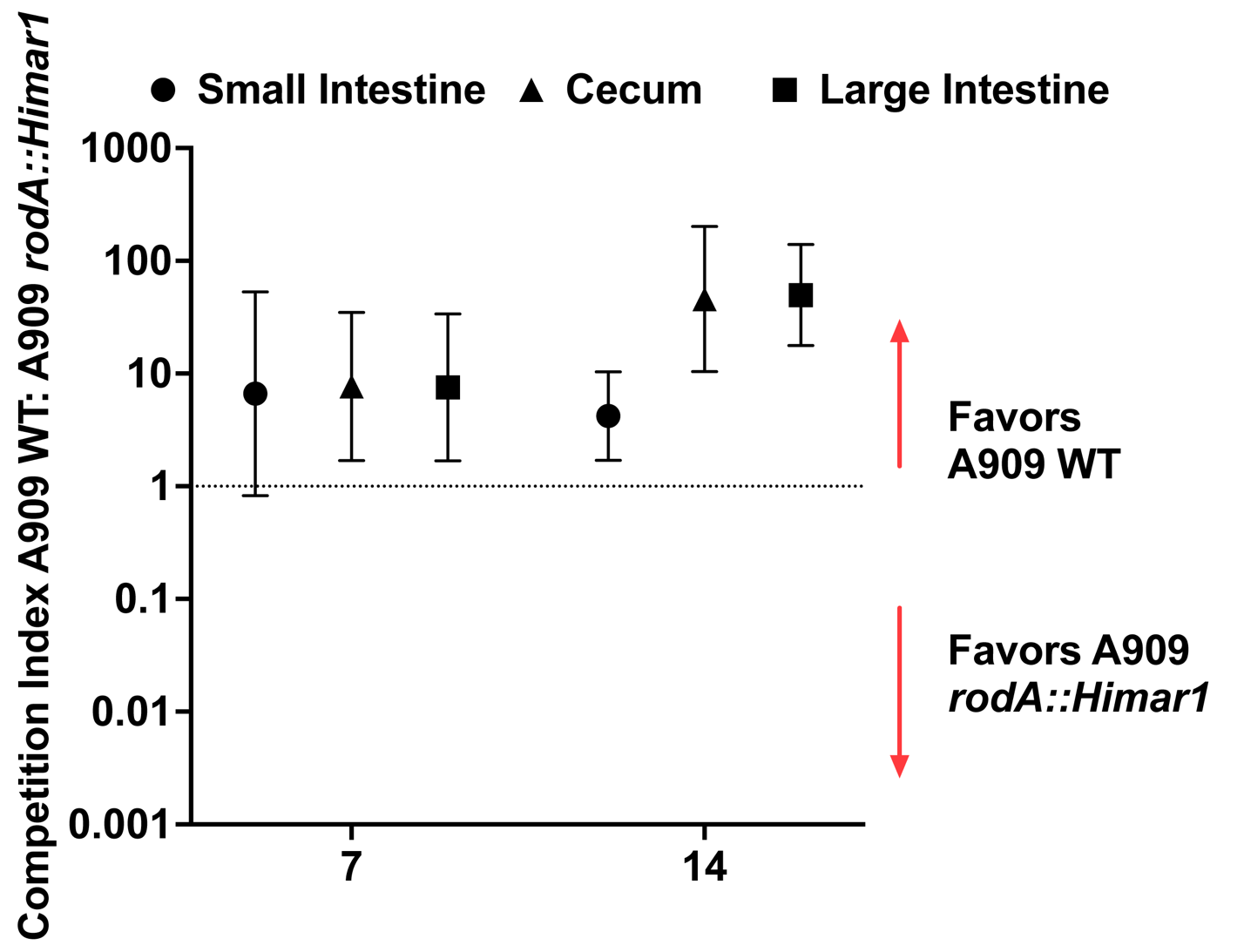

**Days post colonization**

**Supplementary Figure 4.** Genotyping of bacterial colonies from the cocolonization experiment, recovered from the cecum. Representative 10 colonies were analyzed using cPCR, fractionated by electrophoresis in a 1% agarose gel. The left-most lane contains the 1kB DNA ladder, followed by A909 WT control (~3.3kB) and A909Δ*rodA* control (~2kB). The next nine lanes contain representative samples recovered from the cocolonization experiment.

**Supplementary Table 1. PCR Primers used in this study**

| **Name** | **Sequence (5’ to 3’)** | **Notes** |
| --- | --- | --- |
| sgRNA | GGACTAACCCTAATGGTTCTTCC | crRNA sequence (functional RNA component that guides the enzyme to the target DNA). |
| RevCom sgRNA | GGAAGAACCATTAGGGTTAGTCC | Reverse complement to crRNA |
| Fwd Oligo for sgRNA | TAGAT-GGACTAACCCTAATGGTTCTTCC-A | Forward primer for crRNA |
| Rev Oligo for sgRNA | AAATT-GGAAGAACCATTAGGGTTAGTCC-A | Reverse primer for crRNA |
| rodA_Upstream_fwd | aatttctactcttgtagatcGAAGGCTTTGGCAAAATAAC | Homology arms: 780bP sequence upstream (forward) from the gene of interest. |
| rodA_Upstream_rev | cctttttttgAGGACTTCCTTTTTTATTCATG | Homology arms: 780bP sequence upstream (reverse) from the gene of interest. |
| rodA_Downstream_fwd | aggaagtcctCAAAAAAAGGTTGGAAAGATAATCCAGC | Homology arms: 780bP sequence downstream (forward) from the gene of interest. |
| rodA_Downstream_rev | agcttgcatgtctgcaggccCCTGGACGCATGCGAACAG | Homology arms: 780bP sequence downstream (reverse) from the end of the gene. |
| rodA_Chk_fwd | TAGCCATTTTAGGGAGCGCC | Chromosomal Screening Primers to screen and identify mutants: 750 bp upstream from the start gene. |
| rodA_Chk_rvs | TTACCACCGGCATGGAGAAC | Chromosomal Screening Primers to screen and identify mutants: 750 bp downstream of the end of the gene. |
| pGBSComp_rodA_fwd | tgacaatgatgttggatccg  ATGTCACAAAAATCAAACTATTTTAAAG | Forward primer for complementary strain. |
| pGBSComp_rodA_rev | tcgataagcttggctgcagg  TTATATACTACGATACTTCAAATGTG | Reverse primer for complementary strain. |
